# Understanding Author Choices in the Current Conservation Publishing Landscape

**DOI:** 10.1101/2023.08.24.554591

**Authors:** Natalie Yoh, Mukhlish Jamal Musa Holle, Jasmin Willis, Lauren F Rudd, Iain M Fraser, Diogo Verissimo

## Abstract

Conservation literature addresses a broad spectrum of interdisciplinary questions and benefits most by representing a diverse range of authors, particularly those from countries where much conservation work is focused. In other disciplines, it is well known that barriers and biases exist in the academic publishing sphere, which can impact research dissemination and an author’s career development. Here, we used a Discrete Choice Experiment to determine how different journal attributes impact authors’ choices of where to publish in conservation. We identified three demographic groups across 1038 respondents who have previously published in conservation journals, each exhibiting different publishing preferences. Only two attributes showed a consistent response across groups: cost to publish negatively impacted journal choice, including for those in high-income countries, and authors had a consistent preference for double-blind review. Authors from middle-income countries were willing to pay more for society-owned journals, unlike authors from higher-income countries. Journals with a broad geographical scope, which were Open Access, and which had higher impact factors were preferable to two of the three demographic groups. However, we found journal scope and Open Access were more important in dictating journal choice than impact factor. Overall, our findings demonstrate that different demographics experience different preferences or limitations depending on attributes such as a journal’s Open Access policy. However, the scarcity of published authors from low-income countries highlights further, pervasive barriers to representation in conservation research. Based on our findings, we provide recommendations to conservation-related journals to reduce barriers to publishing and ultimately benefit conservation science.

## Introduction

Academic publishing is considered central to the dissemination of scientific research (Medina-Franco & López-López 2022). Academic publications provide a foundation of scientific understanding to inform on-the-ground conservation strategies (Stirling & Burgman 2021). As well as research dissemination, publishing can also be important for a researcher’s career progression. The perceived quality of academic journal publications can affect a researcher’s likelihood of accessing future funding, promotions, and their perceived legitimacy as a researcher (Hall & Page 2015). For researchers based within an organisation such as a non-profit, publishing in reputable journals also increases the visibility of the organisation and can be used to document impact. Therefore, authors must consider how journal choice will ensure the dissemination of their findings, how it will contribute to their careers, and potentially benefit their organisation.

Researchers face multiple considerations and challenges when choosing where to publish, including navigating the many barriers and biases that exist within the publishing sphere. From the author’s perspective, such challenges can be divided into internal and external barriers. Internal barriers may include pressure to conform to Westernised journal styles (Hazen 2016; Prasojo et al. 2019; Oshiro et al. 2020), whereas external barriers may include bias against authors (e.g., geographical or gender discrimination) during the review process, and biases in the perceived value of the research (e.g., journal scope) (Tomkins et al. 2017; Smith et al. 2023). For example, the conservation literature is still considerably biased towards authors from native English-speaking countries, studies focused on vertebrates in terrestrial systems, and positive findings (Di Marco et al. 2017; Stahl et al. 2020; Wood 2020; Amano et al. 2023). While the issues with academic journals have been widely acknowledged across scientific disciplines, the responsibility to overcome barriers has largely been placed on authors rather than on publishers to work towards their removal. For example, many non-native English speakers and non-academic writers are encouraged by journals to seek professional English editing services to better conform to Western scientific styles and written standards (e.g., Hazen 2016 *The International Journal of Logistics Management*). Few conservation journals offer these editing services in-house and even fewer offer these services for free. By requesting authors pay for additional services, publishers are often placing the burden of responsibility on disadvantaged authors to overcome skill barriers rather than work towards equity themselves.

While there has been momentum towards greater inclusivity in conservation research (Raymond et al. 2022), e.g. broadening how journals credit author contributions (Cooke et al. 2022), publishers may still be acting as a bottleneck to the dissemination of conservation knowledge. Few conservation journals have made systemic changes to meet Fair Open Access Alliance standards and authors often face further financial barriers if they wish or are required to make their research publicly available (Veríssimo et al. 2020). Many conservation journals offer a full or partial waiver for these article processing charges (APCs) for authors in low- and middle-income countries. However, critics argue this does not go far enough to address inequity (Rouhi et al. 2022; Sanderson 2023) and Smith et al. (2021) demonstrated waivers were ineffective at increasing the representation of low-income authors. Such ineffectiveness may relate to poor communication surrounding waiver eligibility, waiver restrictions, or - less often discussed - negative perceptions towards waivers, which may be seen as patronizing by those they aim to help (Meagher 2021; cOAlition S 2024). Others argue that APCs pose a barrier to many more authors who do not qualify for waivers, including authors from high-income countries based at smaller academic institutions or in non-academic settings (Frank et al. 2023; Byrne 2024).

Understanding to what extent journal characteristics, such as APCs, factor into journal choice and how this varies across author demographics and psychographics (psychological and behavioural variables) can therefore inform journal reform and facilitate more inclusive learning and knowledge exchange. In this study, we assess preferences for certain journal attributes for researchers who have published in conservation science. Specifically, we assess the interplay between seven different journal attributes and how they impact an author’s journal choice. Here, we focus on published authors to gauge preferences within the current publishing landscape. Hence, we are not investigating hard barriers that prevent *potential* authors from publishing. Rather, we are interested in how these attributes impact choice for those already active within the publishing system. We subsequently contextualise the impact of current publishing decisions in conservation research and provide recommendations to conservation-related journals on how to reduce challenges to publishing.

## Methods

### Survey design

Our questionnaire consisted of (1) a brief description of the survey background, (2) questions related to respondents’ demographics, (3) a Discrete Choice Experiment focused on seven key journal attributes, and (4) a section for the respondent to rank conservation journal attributes (Appendix 1). In the first section, we confirmed whether the respondents had previously published in a peer reviewed, conservation-related journal and if so, how many conservation-related papers they had published within the past year (any author position). In the second section, we collected the respondent’s demographic information including age, nationality, country of residence, and racial identity. Providing such information was voluntary throughout the survey. We chose to collect demographic information at this stage in the survey to collect information about non- or incomplete survey respondents. However, we acknowledge that asking demographic questions at this stage in the survey may prime authors to reflect on their demographic characteristics while completing the remainder of the survey (Hughes et al. 2016). We determined which journal attributes to include in the Discrete Choice Experiment following a workshop and online questionnaire where we asked attendants at the International Conference of Conservation Biology (ICCB), 2021 about how they choose where to publish (Appendix 2). Following this preliminary data collection, we identified seven main attributes that informed researchers publishing decisions (Figure 1). We used these seven attributes to generate a Discrete Choice Experiment using an orthogonal design generated in IBM SPSS 22.0 with the initial choice alternatives coupled using a “shifted technique” (Louviere et al. 2000) into 16 trichotomous choices. The “shifted technique” introduces variation in the attribute levels between choice sets to better capture how decisions can vary depending on differences in the scenarios presented. We provided an opt-out choice in the form of “Would not choose any of these journals” (Choice [d]). Within our study design, we referenced existing conservation and ecology journals to ensure attribute values were reflective of journals in the discipline.

**Figure 1.**
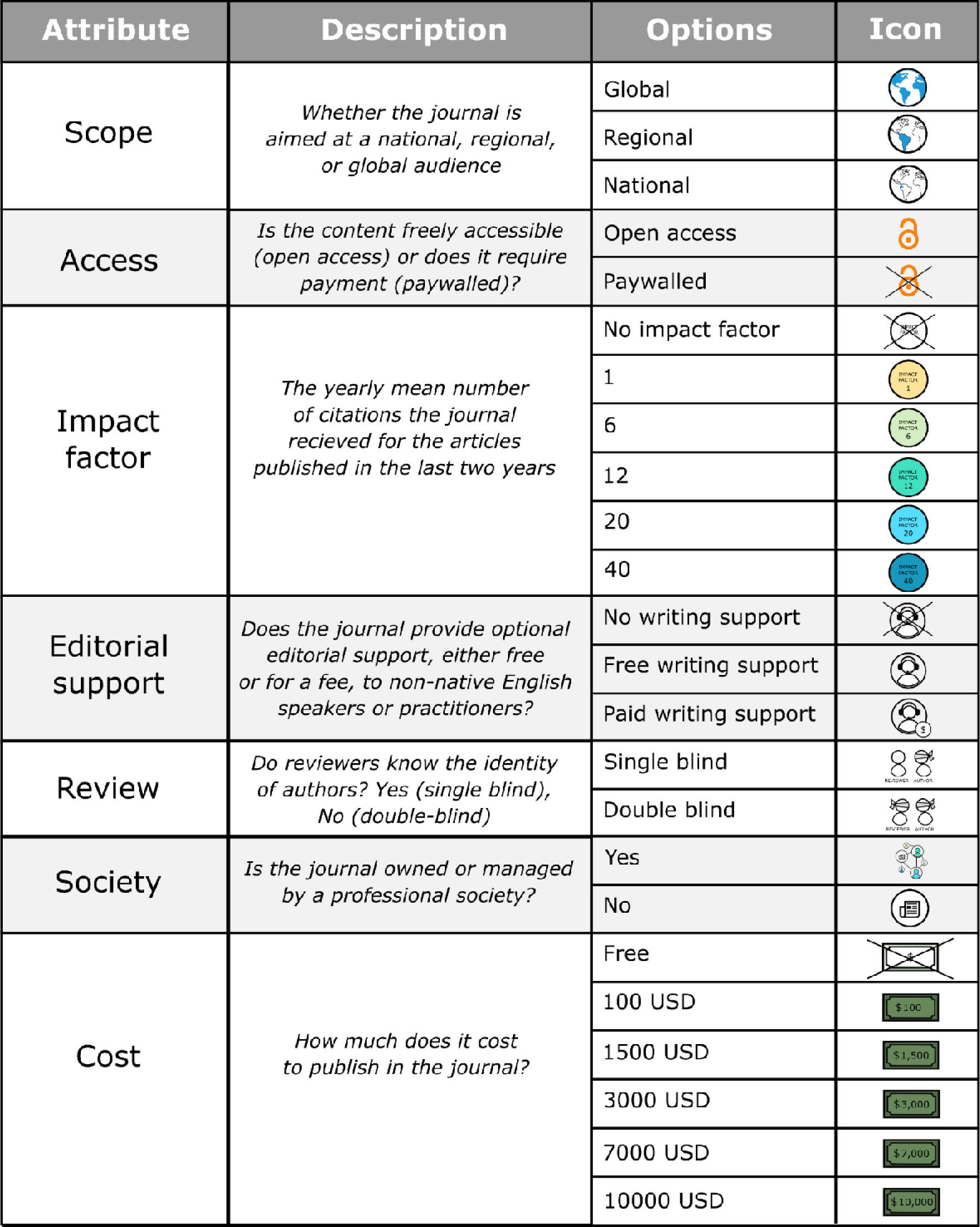
Attributes and levels of the Discrete Choice Experiment investigating journal preference among conservation authors.

The resultant data were analysed using a multinomial logit model and parameter estimates of the main effects were used as priors in a D-efficient Bayesian design implemented in Ngene 1.0.1 to design the final choice sets. We chose a multinomial logit model to analyse the resultant data as we asked respondents to choose between three choices or the opt-out choice. We specified a D-efficient Bayesian prior design, where the parameter is described by its probability distribution rather than a fixed value, to optimize the experiment design and improve precision (Rose & Bliemer 2009). Parameter estimates of the main effects within the multinomial logit model were used as the priors in this design. Using 500 Halton draws from normal prior distributions for each parameter, we compared the mean Bayesian D-error of over 50,000 designs. The Bayesian D-error quantifies how good or bad a design is at extracting information from respondents in an experiment. We selected the model with the lowest error at 0.1606. We limited the number of scenarios to twelve to keep the Discrete Choice Experiment design simple and to limit respondents’ cognitive burden. In the last section, we asked the respondents to rank the attributes from the most to the least prioritised.

### Data collection

We distributed the survey in two ways, (1) directly through authors’ email addresses that we collected from published conservation articles, and (2) indirectly via communication platforms of conservation-related institutions and organisations (newsletters, mailing lists, and social media platforms; Appendix 3). Email addresses were collated for all authors (including corresponding and non-corresponding authors) who published in 18 conservation-related journals within 2010 and 2020 (Appendix 3), where contact information could be publicly obtained from recent publications or their affiliation webpage (n = 9,994). We collected the data using SmartSurvey premium (www.smartsurvey.co.uk), an online survey software and questionnaire tool, between 19 August and 3 November 2022. We offered respondents the opportunity to enter a raffle with the chance to win three 1-year memberships and three 3-year memberships for the SCB as incentives for completing the survey. Only respondents who had published an article in a peer reviewed journal were considered in the analysis. This project has been reviewed by and received ethics clearance from RETRACTED FOR REVIEW.

### Data analysis

We used different modelling approaches to investigate preferences for (a) all respondents, and (b) distinct subpopulations. First, we used a multinomial logit model to evaluate the preferences of the entire sample of respondents. We used dummy coding to represent categorical variables with more than two categories (e.g. Journal Scope) in the model estimation (Appendix 4). This method assumes homogenous preferences across respondents. As different author demographics may demonstrate different preferences between attributes, we were also interested in grouping respondents into distinct subpopulations observed in the total sampled population. To investigate this potential heterogeneity in choice, we employed two model specifications. First, we estimated a latent class model which allows for homogeneous finite classes. Each class represents a distinct segment of the respondent population, hereafter referred to as a segment (Boxall & Adamowicz 2002). After a careful examination of the data, segment membership was explained by the inclusion of three socio-economic variables: a dummy variable for high or lower income (High Income); age of respondents (Age); and the number of publications (Number of Publications). High or lower income were defined by the World Bank income groups (2022), where the latter is defined as both middle-income and low-income countries. However, due to the representation of respondents, lower income predominantly represents those from middle-income countries. The appropriate number of segments (e.g. the number of distinct subpopulations observed) was determined by examining a range of model statistics, including Akaike information criterion (AIC) and Bayesian information criterion (BIC). Second, we estimated a mixed logit model in Willingness to Pay space assuming all attributes follow a normal distribution (Balcombe et al., 2010). For all models to capture the "neither" responses, we included an alternative specific constant (ASC). When "neither" was selected, ASC assumed a value of one, showing the utility gained from not selecting any of the available choice options. Both models are explained in detail in Appendix 5. Model comparison statistics (i.e. AIC, BIC) have been generated for all model specifications examined (Appendix 6). Although the mixed logit model performed well, we focus our attention on the latent class model as it yields greater insights into respondent preferences. Full results for the mixed logit are reported in Appendices 7 and 8.

For the latent class models, we found that as we increased the number of segments from three to four, the results started to become behaviourally unrealistic and unstable regarding the magnitude of the implied Willingness to Pay estimates. Thus, a three-segment latent class model is the model we report. With this model and the multinomial logit model (and for the mixed logit, see Appendix 7) we report Willingness to Pay for all attributes. Willingness to Pay is the maximum price in US dollars a respondent is willing to pay for a particular attribute, such as a higher impact factor. For the multinomial logit model and the latent class model, we generated standard error estimates using the Wald method.

## Results

A total of 1531 people responded to the survey between 19 August to 3 November 2022. Of these, 1199 respondents completed the survey, with 1038 respondents (86.57%) reported to have published a conservation-related study in a peer reviewed journal. On average, respondents had published a mean of 3.28 papers over the previous year (SD ± 5.26) and were 40 years old (SD ± 11.31) (Appendix 9). Most respondents were from the USA (165 respondents; 15.90%), India (110 respondents; 10.60%), and the UK (84 respondents; 8.09%). Approximately half of the respondents (483 respondents, 46.53%) identified themselves as White Europeans/North Americans/Australians/New Zealanders, 12.04% as South Asian, 8.67% as Southeast Asian, 7.61% as Latino/Latina/Latinx, and 7.03% as Black African. In total, we obtained 12,365 choice cards from 1038 respondents. Table 1 presents the main model results for the multinomial logit and the three-segment latent class model. For the latent class model, we also include the segment membership functions.

**Table 1.**
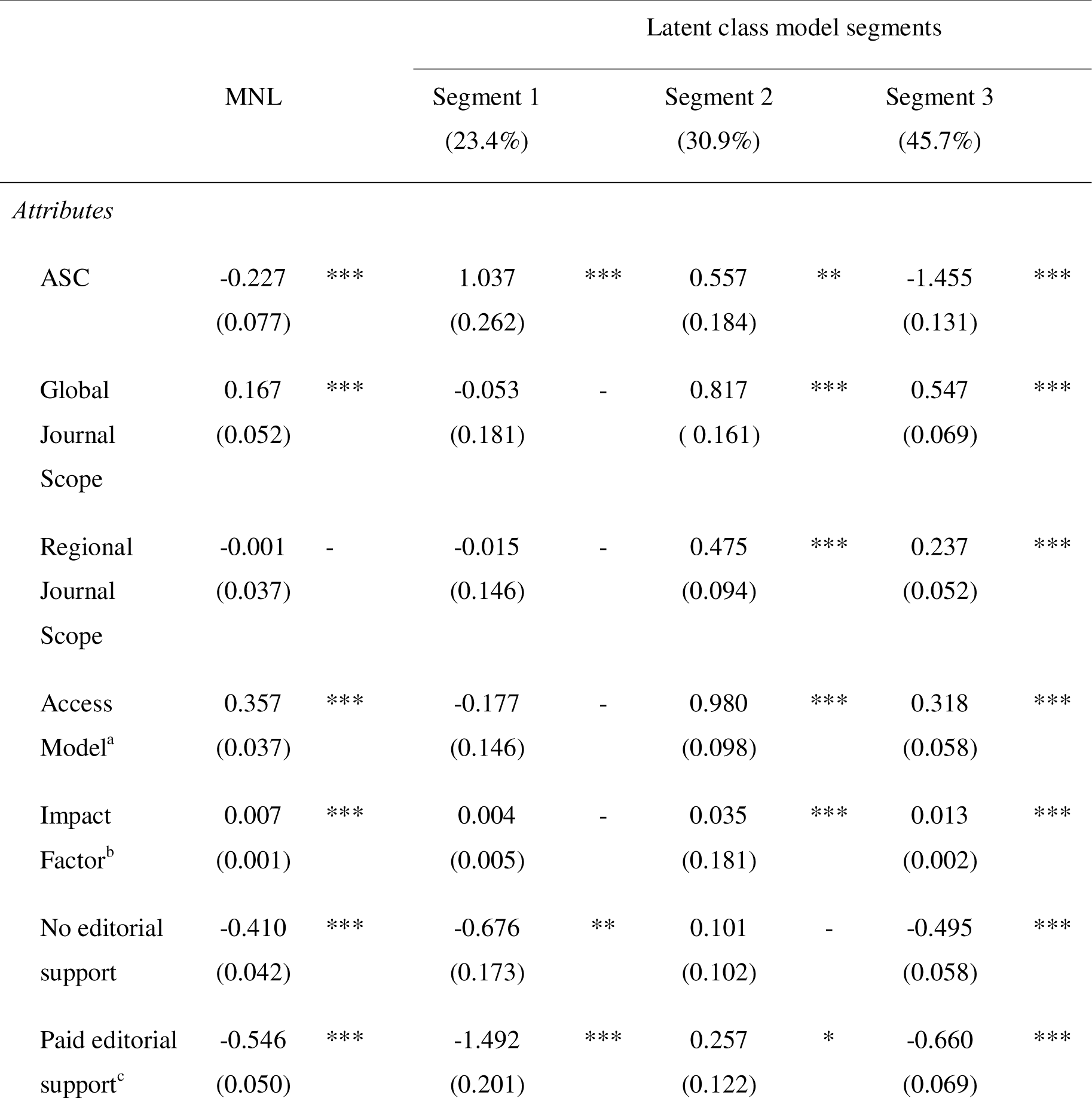
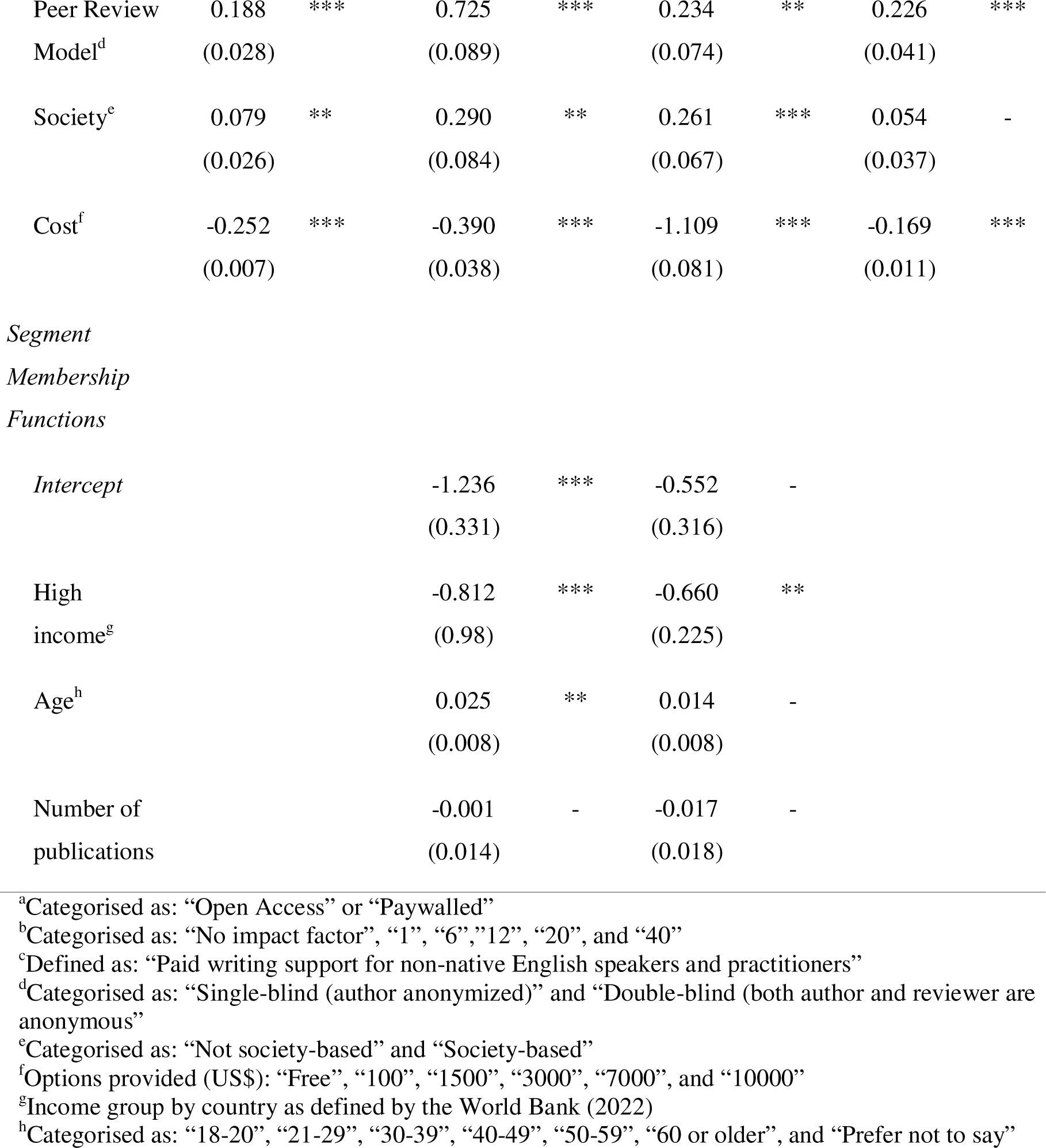
The multinomial logit (MNL) and latent class model (LCM) estimates of utility function for each attribute, including standard errors (in brackets). Significance levels: \**P* < 0.05, \*\**P* < 0.01, *** *P <* 0.001. ASC – Alternative specific constant. Segment 3, as the reference class, represents younger respondents from high-income countries. Segment 2 represents more respondents from lower income countries. Segment 1 represents more respondents from lower income countries and older respondents.

When asked to rank the attributes, respondents chose: (1) journal scope, (2) whether a journal was Open Access, (3) impact factor, (4) cost, (5) options for editorial support, (6) whether a journal offered double-blind review, and (7) whether the journal was associated with a society, from the most important to the least important. Although Peer Review Model was ranked low, Willingness to Pay suggests it is important in journal choice (Appendix 7). A total of 312 respondents (30.65%) stated that they had ignored attributes.

For both models reported (Table 1), we can see that the cost attribute is negative and always statistically significant. Turning to Segment 1, this segment represents 23.4% of respondents, and they are older and from lower income countries given the segment membership function estimates. Lower income here includes low-income countries but predominantly represents middle-income countries due to a lack of respondents from low-income countries. The most statistically significant attributes for segment 1 were the availability of editorial support (demonstrating a preference for free support over no support or paid support), a preference for journals offering double-blind review, and journals which are associated with a society. For Segment 2, the second largest group of respondents (30.9%) we find that more of the attributes are statistically significant compared to Segment 1. In particular, Segment 2 demonstrated a preference for journals with a global or regional scope, Open Access, and journals with higher impact factors. Segment 2 also responded positively to paid editorial support, unlike other segments. Given the segment membership function estimates for Segment 2, respondents appear to be younger authors from lower income countries. Segment 3 contains the largest proportion of respondents (45.7%) with all attributes being statistically significant except whether a journal is associated with a society. Segment 3 demonstrated similar preferences to Segment 2 except for editorial support. Segment 3 avoided journals with no support or paid editorial support, comparable to Segment 1. The class membership function is not estimated for Segment 3 given the model identification restrictions. However, we can infer that respondents likely to be in this segment will be younger (the sign for this parameter is positive for both Segments 1 and 2), from higher-income countries (the sign for this parameter is negative for both Segments 1 and 2) and with higher numbers of publications (the sign for this parameter is negative for both Segments 1 and 2). The results from the mixed logit in WTP for our attributes (see Appendix 8) reveal that WTP is positively related to income, negatively related to age and negative but statistically weak for the number of papers published. We present Willingness to Pay estimates for both the multinomial model and the latent class model (Table 2).

**Table 2.**
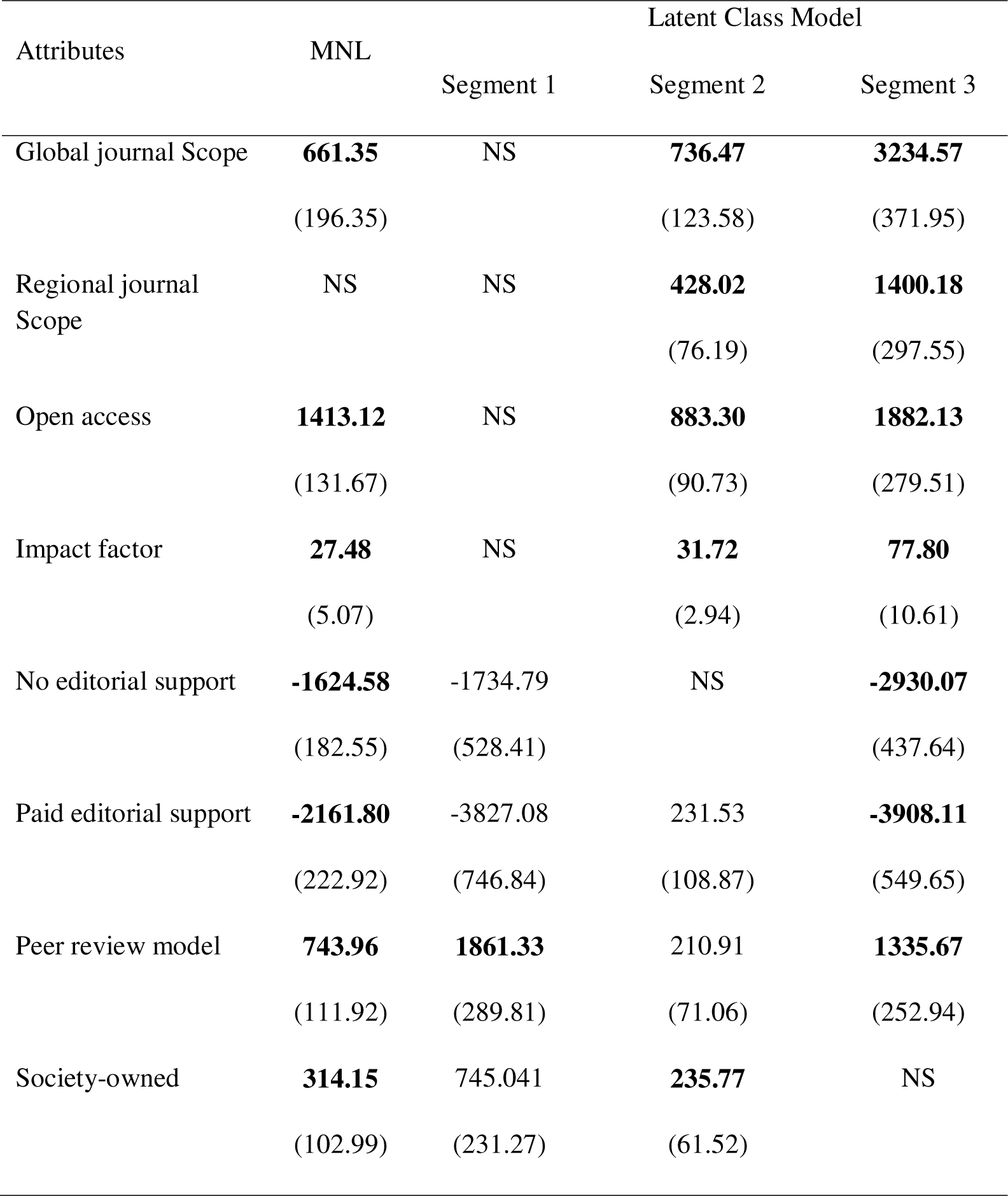
Willingness to Pay for different attributes of academic journals, only statistically significant values in terms of influencing choice are provided (p < 0.05). Bold represents values with significance < 0.01. All values are provided in US dollars. Values in brackets are standard errors generated using the Wald method. NS – not statistically significant.

| Attributes | MNL | Latent Class Model |  |  |
| --- | --- | --- | --- | --- |
|  |  | Segment 1 | Segment 2 | Segment 3 |
| Global journal Scope | **661.35**<br>(196.35) | NS | **736.47**<br>(123.58) | **3234.57**<br>(371.95) |
| Regional journal Scope | NS | NS | **428.02**<br>(76.19) | **1400.18**<br>(297.55) |
| Open access | **1413.12**<br>(131.67) | NS | **883.30**<br>(90.73) | **1882.13**<br>(279.51) |
| Impact factor | **27.48**<br>(5.07) | NS | **31.72**<br>(2.94) | **77.80**<br>(10.61) |
| No editorial support | **-1624.58**<br>(182.55) | -1734.79<br>(528.41) | NS | **-2930.07**<br>(437.64) |
| Paid editorial support | **-2161.80**<br>(222.92) | -3827.08<br>(746.84) | 231.53<br>(108.87) | **-3908.11**<br>(549.65) |
| Peer review model | **743.96**<br>(111.92) | **1861.33**<br>(289.81) | 210.91<br>(71.06) | **1335.67**<br>(252.94) |
| Society-owned | **314.15**<br>(102.99) | 745.041<br>(231.27) | **235.77**<br>(61.52) | NS |

We find variation between the models in terms of the magnitude of Willingness to Pay as well as the signs of the estimates (Table 2). We find that for all attributes the multinomial logit and Segment 3 of the latent class model are the same signs. However, Segment 3 yields by far the largest Willingness to Pay estimates compared to all other model results. There are some similar results between Segments 2 and 3 which showed a preference for a broad journal scope, Open Access, and higher impact factors. However, the estimates for Segment 3 are significantly bigger than for Segment 2: global or regional journal scope ($3235 and $1400 respectively), Open Access ($1882 and $883), and higher impact factors ($78 and $32). None of these attributes are statistically significant for Segment 1.

Peer Review Model was important for all segments, each yielding a positive Willingness to Pay estimate, and in this case, Segment 1 yielded the largest estimate of $1861 compared to $1336 for Segment 3. Both Segments 1 and 2 were also willing to pay for journals that were society-owned, especially Segment 1. We see variation in the Willingness to Pay estimates for editorial support between segments, where Segment 2 showed a slight preference for journals offering paid editorial support. However, both Segments 1 and 3 expressed negative preferences for both no- and paid-editorial support options. Finally, when we compare the magnitude of our Willingness to Pay estimates with those of the mixed logit (see Appendix 6), we see that the mixed logit yields on average lower estimates. The signs of the estimates in Appendix 7 are consistent with those reported in Table 2, but only journal impact factor ($14) and peer review ($795) now yield estimates that are similar in magnitude. These results indicate that we need to treat our latent class model Willingness to Pay estimates as being on the high side.

## Discussion

Overall, we demonstrated publishing preferences within conservation are not homogenous across published authors. Cost and Peer Review Model were the only two attributes to which all respondent segments responded consistently. Furthermore, we show that preferences for different journal attributes differ between World Bank income groups. These results highlight how different attributes may act as filters for different author demographics. We also demonstrate how journal preference is multi-faceted as no one factor dictated journal choice. Impact factor is often touted as a major driver of journal choice (Nicholas et al. 2014). However, despite ranking in the top three most influential attributes when self-reporting, we found respondents were willing to pay little for higher impact factors compared to other attributes, such as a global/regional scope or for Open Access. Unlike impact factor, many important attributes dictating journal choice are governed by the publisher. Therefore, publishers have the capacity to attract more authors and acknowledge their responsibility to uphold equitable publishing opportunities in conservation journals.

### Reducing the cost of access

Open Science, including free-to-read publishing, represents a positive step towards greater transparency and accessibility to scientific knowledge. Currently, however, Open Access represents a double-edged sword for inclusion. While Open Access mandates ensure researchers are obligated to make their research available to all, APC Open Access models can prevent research from being published at all for those who lack the resources to pay these fees. Much of the discussion regarding the “APC-barrier” (Klebel & Ross-Hellauer 2023) has focused on authors in low- and middle-income countries, as they are likely to be the most restricted by APCs. However, as our results demonstrate, lower costs are preferential to authors across all demographics, including those in high-income countries. Of the 18 conservation-related journals we used to collate author email addresses, APCs ranged from $1632 (or $2040 for non-society members) to $4600 (Appendix 10). Transformative agreements, including Read and Publish agreements (financial agreements between academic institutions and publishers whereby researchers can publish Open Access without charges), aim to shift the financial burden of publishing from authors onto institutions. However, currently, these agreements reinforce existing inequities. At their best, Read and Publish agreements can restrict publishing options for authors by limiting which journals the institution will cover. At their worst, Read and Publish agreements limit who can contribute to the scientific conversation to only the most well-funded, most-resourced academic researchers (Debat & Babini 2019). Neither these agreements nor waivers aim to change the status quo of commercial publishing models which prioritise profits over scientific dissemination (Debat & Babini 2019; Byrne 2024). Our findings demonstrate that substantially reducing or, preferentially, removing costs would benefit all authors.

Nevertheless, hosting a journal requires infrastructure. Many societies outsource this requirement to commercial publishers, who in turn can help generate funds for the society to support activities such as conferences, education/training, and future research, through publication fees. As such, many learned societies rely on their journal portfolio as vital revenue streams (Fyfe et al. 2017; Fyfe 2023). However, this is at odds with many academic societies’ commitments to Diversity, Equity, and Inclusion where publication fees act as a barrier to the very authors the society aims to support. Academic societies committed to addressing inequities in publishing in conservation may consider converting their journals to diamond Open Access models, such as with *Edinburgh Journal of Botany,* or to initiatives such as *Peer Community in [Ecology]* and *SciPost* (PCI Ecology 2023; SciPost Foundation 2024). These models aim to provide a free alternative to traditional Open Access model journals.

*Edinburgh Journal of Botany* is published by the Royal Botanic Garden Edinburgh, a charity and non-departmental UK Public Body. The journal was established in 1900 and moved to a diamond Open Access model in 2021 (RBGE 2024). Similarly, *Chemical Science* is a diamond Open Access journal for the Royal Society of Chemistry which received 6980 submissions in 2023 and has a 5-year impact factor of 8.6 (RSC 2024). The journal has options for both single and double-blind review, fast processing times, and optional transparent peer review (e.g. where the reviewer comments and responses are published alongside the final article; RSC 2024). Both represent different pathways for conservation societies to adopt free-to-publish-free-to-read publishing models. In doing so, societies can ensure it is the quality of the peer review, not the price to publish, that dictates our perception of quality in conservation literature.

Currently, most society-owned diamond Open Access journals rely on society membership fees to cover operational costs (Bosman et al. 2021). We found younger respondents in high-income countries were less likely to prioritise society-owned journals. As early as 2008, concerns were raised over the decline of young professionals joining academic societies, such as the SCB (Schwartz et al. 2008; Grajal 2009). Grajal (2009) argued academic societies need to explore ways to increase their value for younger conservation professionals and our research indicates an opportunity to do so specifically in the publishing domain.

### Promoting equitable peer review

Several studies have demonstrated how single-blind reviews can offer advantages to authors from high-income, English-speaking countries across biological science journals, including in *Functional Ecology,* which is likely due to prestige bias (Fox et al. 2023; Smith et al. 2023) (e.g., where reviewers expect work from certain countries, institutions, or individuals to be of higher quality). Despite this, Smith et al. (2023) found only 15.9% of 541 biological science journals practised double-blind review, including universal double-blind review (e.g., *Conservation Biology*) or optional double-blind review (e.g., *Nature Ecology and Evolution*). We found all author segments demonstrated a preference for double-blind over single-blind peer review. Previous research in the medical literature (Parmanne et al. 2023) has also shown transitioning to double-blind review does not impact a reviewer’s willingness to review. Therefore, introducing double-blind review is a cost-neutral way for journals to attract new authors, while at the same time working towards reducing unconscious (or conscious) bias in peer review.

Anonymising work merely represents an initial step in mitigating bias within the peer review process. Anonymised authors may still face discrimination from reviewers and editors should they diverge from predetermined language and style criteria, irrespective of the scientific merit of their work. Nevertheless, despite an increased likelihood of journal rejections, Amano et al. (2023) found many authors do not or cannot access paid, professional editing services, particularly those from low and middle-income countries. Our results indicate that two of the three segments actively avoid journals with the option to pay more for editorial support, even when compared to journals that offer no support. More concerning still is the insistence that authors specifically seek collaborations with native-English speakers to ensure they produce “high-quality research articles” (Balan 2021). Needless to say, collaboration should be born from more than someone’s first language and such comments are an insult to the groundbreaking work undertaken by non-native English speakers around the world. If journals insist that publications are written in English (but see Amano et al. 2021, 2023; Chowdhury et al. 2022), it should be their responsibility to support authors, not merely the burden of non-native English-speaking authors. Offering such assistance would in return aid reviewers, whose voluntary role and expertise should be to assess the quality of the research. *Conservation Biology* offers an alternative strategy to support authors through their Publication Partner Program (SCB 2023). This free initiative invites authors to partner with an experienced volunteer who can help with manuscript revisions, aiming to improve the likelihood of publication. Such peer-support strategies help to acknowledge systemic barriers and provide training and support to those who are disadvantaged by the current publishing environment. However, it is important to note that we found certain respondents did not exhibit a preference for free support over no editorial support. Thus, more research is needed to ensure support schemes are meeting the needs of those they aim to help.

### Who is missing from the conversation?

We found that all segments published similar numbers of publications over the last year, suggesting journal preferences were not affecting how many publications our respondents were able to publish among segments. Yet, given that just three countries represented over a third of our respondents and almost half of the respondents identified as White Europeans/North Americans/Australians/New Zealanders, our sample suggests barriers are hindering a more diverse representation of authors from publishing in the first instance. Our sample was largely dominated by respondents from high-income and upper-middle-income countries (51.8% and 20.1% of respondents respectively) with few respondents from low-income countries (2.0%) (Appendix 9). Consequently, our sample does not aim to capture hard barriers to publishing faced by authors in lower-middle and low-income countries, be it related to costs, review bias, or other factors. Subsequent research should survey both current authors and potential authors to understand the publishing preferences and/or barriers more comprehensively for authors in these regions. This may include assessing differences in how Willingness to Pay is expressed differently across different demographics (i.e. cannot pay versus choose not to pay). Future research could also explore how other dimensions of an author’s identity affect journal choices, such as gender, discipline (e.g., social science vs natural science vs humanities), industry (e.g., between academia and non-academic sectors), focal taxa or ecosystem, career stage and tenure (Griffiths & Dos Santos 2012; Teel et al. 2018; Maas et al. 2021). There may also be additional journal attributes, such as perceptions of peer review quality, that were not captured by the initial workshop and questionnaire at ICCB. Such attributes may be associated with certain demographics that were not well represented by those at ICCB, such as peer review speed for early-career researchers (Nguyen et al. 2015). Future research could explore how choice is affected by measurable or perceived differences in the review process among journals.

### How perceptions of research value impact conservation

Although regional or local scale research is often most informative for on-the-ground conservation practitioners (Stergiou & Tsikliras 2006; Calver et al. 2010), academic career incentives do not necessarily align with impact outside of academia (Rigby et al. 2015). Authorship in journals considered to be high-ranking or have high prestige is perceived as important for a researcher’s career progression (Rigby et al. 2015; Nicholas et al. 2017) and many of the most highly ranked conservation journals now prioritise studies with a broad geographical or taxonomical scope. These large-scale studies are often beneficial for identifying priority areas of future research or intervention. Cultivating research deemed to be applicable for a broader audience is also beneficial to the publisher as it increases the likelihood of citations; the currency used to bolster the perceived importance and legitimacy of a journal, and therefore the likelihood of future submissions. This perpetuates a cycle of science that is not necessarily engaged with applied conservation needs. In this way, publishers run the risk of creating a dilemma for researchers – a trade-off between safeguarding their career and securing funding versus contributing to local conservation efforts. Learned societies have a unique opportunity to capitalise on their authority and credibility to challenge this perception of value. Unlike new independent journals, journals run by learned societies do not rely on impact factor, and thus citations, to demonstrate their scientific legitimacy. Free of this constraint, such journals have more freedom to find new ways to recognise meaningful contributions to science beyond the number of citations. For example, providing publishing opportunities that prioritize science that informs conservation impact (but potentially within a limited scope) over highly citable, global reviews with limited conservation applications. While both are valuable contributions to the academic literature, there are many publishing opportunities for the latter but few prestigious journal portfolios offer the former (but see *Conservation Science and Practice*; SCB 2024).

### Conclusions

Here, we have outlined several ways in which journals themselves can impact the conservation literature and future research directions. Given our findings, we suggest several recommendations for publishers that would promote better equity, diversity, and inclusion in conservation publishing and ultimately benefit conservation science. We recognise systemic issues in academic publishing go beyond the conservation literature. In all fields, commercial publishers lack incentives to deviate from a publishing landscape that fails authors and readers when they seek to benefit from the substantial profits it generates (Van Noorden 2013). In a time when the market is becoming increasingly saturated by predatory publishing practices, learned societies are presented with the opportunity to overhaul how we disseminate and value science. Conservation is distinct from other sciences due to its interdisciplinary nature, its close ties to policy and management, and the need to be dynamic in the face of changing environmental conditions and public perceptions. Therefore, conservation literature benefits more than most by having the largest diversity of voices at the table. Finally, we also acknowledge that publishing is the last stage in the research pipeline. Many people will have already been excluded from the publishing process in the research planning and execution stages. Collectively, we are all responsible for improving equity, diversity, and inclusion across the whole research timeline.

## Supporting Information

Additional supporting information may be found in the online version of the article at the publisher’s website

---

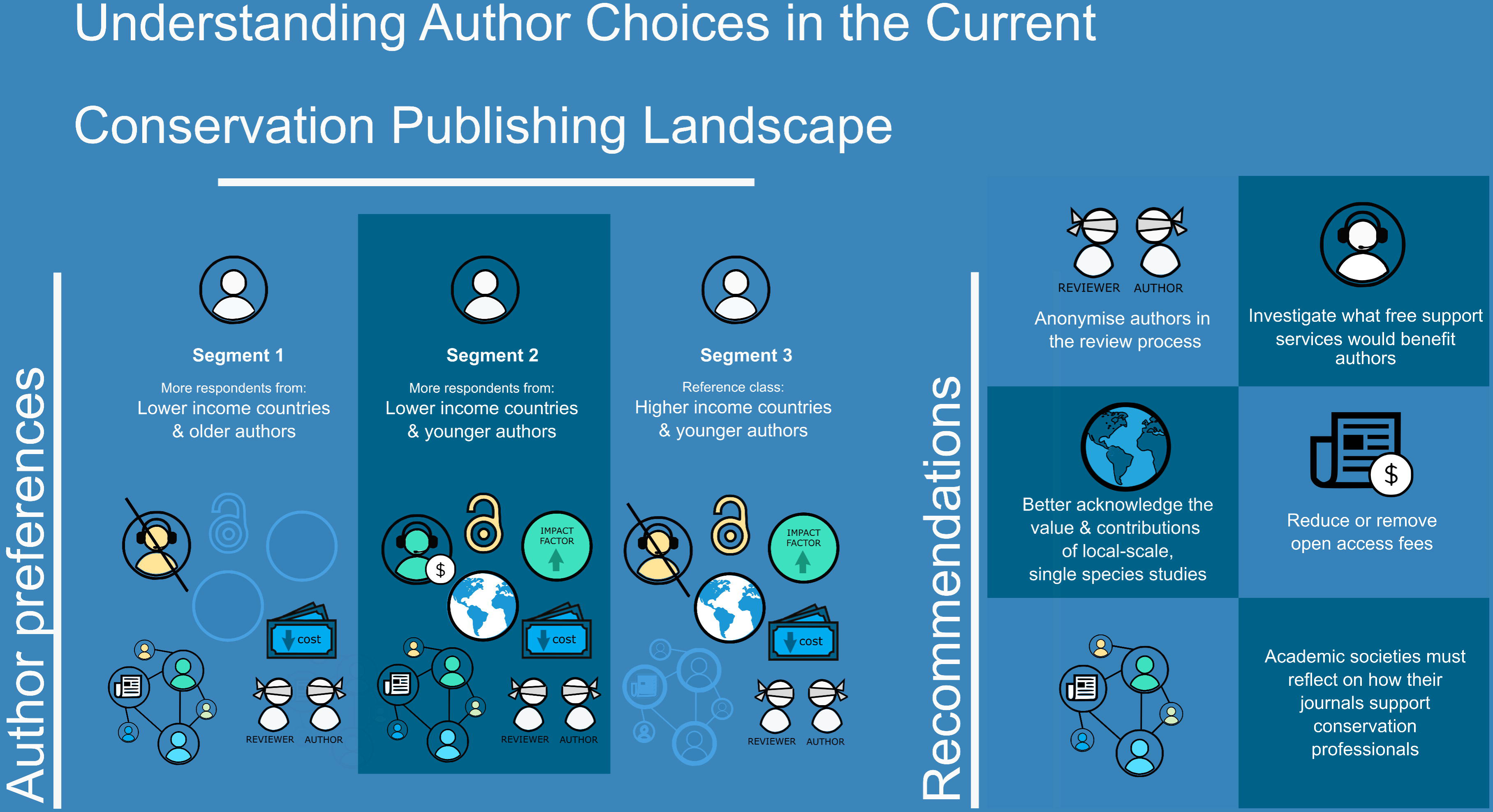

