## Supporting Information for "Understanding Author Choices in the Current Conservation Publishing Landscape"

**Appendix 1.** Example of the survey including the background information provided to respondents of the Discrete Choice Experiment

**Background information**

1. What is the aim of this study?

You have been invited to complete the survey below as a conservation researcher. The study aims to identify the preferences of different groups (e.g. social or ethnic groups) when choosing conservation research publishers, in order to identify strategies to improve inclusivity in conservation publishing. This study is supported by the Society for Conservation Biology (SCB). The Principal Researcher is RETRACTED FOR REVIEW, who is attached to the RETRACTED FOR REVIEW as a RETRACTED FOR REVIEW.

2. What do I need to do?

You are being asked to complete a demographic survey and to respond to several choices about conservation publication. The survey should take approximately 10 minutes to complete. Please note that the survey responses are anonymous. By submitting your response to this survey, you are consenting that your data can be anonymously used for the study.

3. Do I have to take part?

No. Please note that participation is voluntary. If you do decide to take part, you may withdraw at any point for any reason before submitting your answers by pressing the ‘Exit’ button/closing the browser.

4. How will my data be used?

All data will be kept in a password-protected file within the RETRACTED FOR REVIEW secure network and will not be shared with any third parties. We will keep all data records from the survey responses for three years post the publication of this research. Your contact details were obtained from publicly available sources, but will be deleted when the study has been completed.

5. Who has reviewed this study?

This project has been reviewed by, and received ethics clearance through, a subcommittee of the RETRACTED FOR REVIEW.

6. Who do I contact if I have a concern or I wish to complain?

If you have a concern about any aspect of this study, please speak to RETRACTED FOR REVIEW (RETRACTED FOR REVIEW), and we will do our best to answer your query. We will acknowledge your concern within 10 working days and give you an indication of how it will be dealt with. If you remain unhappy or wish to make a formal complaint, please contact the RETRACTED FOR REVIEW who will seek to resolve the matter as soon as possible: Email: RETRACTED FOR REVIEW; Address: RETRACTED FOR REVIEW.

Please contact RETRACTED FOR REVIEW (RETRACTED FOR REVIEW) for more information about this study.

**Demographic survey**

1. If you have read the information above and agree to participate with the understanding that the data you submit will be processed accordingly, please tick the box below to start. *

   ☐ Yes, I agree to take part.
2. Please note that you may only participate in this survey if you are 18 years of age or over. *

   ☐ I certify that I am 18 years of age or over.
3. Have you ever published a conservation-related study in a peer-reviewed journal? *

   ☐ Yes

   ☐ No
4. If you have published a conservation-related paper in a peer-reviewed journal, how many papers did you publish last year? (*Fill with number*)
5. What is your age? *

   ☐ 18-20

   ☐ 21-29

   ☐ 30-39

   ☐ 40-49

   ☐ 50-59

   ☐ 60 or older

   ☐ Prefer not to say
6. What is your nationality? The question allows for multiple options to be selected *

   *Tick box options including “Prefer not to say” and “Unknown”.*
7. What is your country of residence * [Dropdown menu]

   *Dropdown menu including “Prefer not to say”.*
8. How would you best describe your race? This question allows for multiple options to be selected *

*Tick box options including "No response" and "Other option not listed here".*

### Introduction to Choice Experiment

You will be now asked to make a series of choices between academic journals with different characteristics. Please answer as if you were considering where to publish your next paper. These characteristics are:

| Attribute | Description | Options | Icon |
|---|---|---|---|
| Scope | *Whether the journal is aimed at a national, regional, or global audience* | Global | |
| | | Regional | |
| | | National | |
| Access | *Is the content freely accessible (open access) or does it require payment (paywalled)?* | Open access | |
| | | Paywalled | |
| Impact factor | *The yearly mean number of citations the journal recieved for the articles published in the last two years* | No impact factor | |
| | | 1 | |
| | | 6 | |
| | | 12 | |
| | | 20 | |
| | | 40 | |
| Editorial support | *Does the journal provide optional editorial support, either free or for a fee, to non-native English speakers or practitioners?* | No writing support | |
| | | Free writing support | |
| | | Paid writing support | |
| Review | *Do reviewers know the identity of authors? Yes (single blind), No (double-blind)* | Single blind | |
| | | Double blind | |
| Society | *Is the journal owned or managed by a professional society?* | Yes | |
| | | No | |
| Cost | *How much does it cost to publish in the journal?* | Free | |
| | | 100 USD | |
| | | 1500 USD | |
| | | 3000 USD | |
| | | 7000 USD | |
| | | 10000 USD | |

If none of the options provided is acceptable, you also have the option to not choose.

### Choices

9-20. Imagine you have a manuscript ready for publication and you are about to submit your manuscript. There are three options of journal publishers with different services as follows; which alternative would you choose?

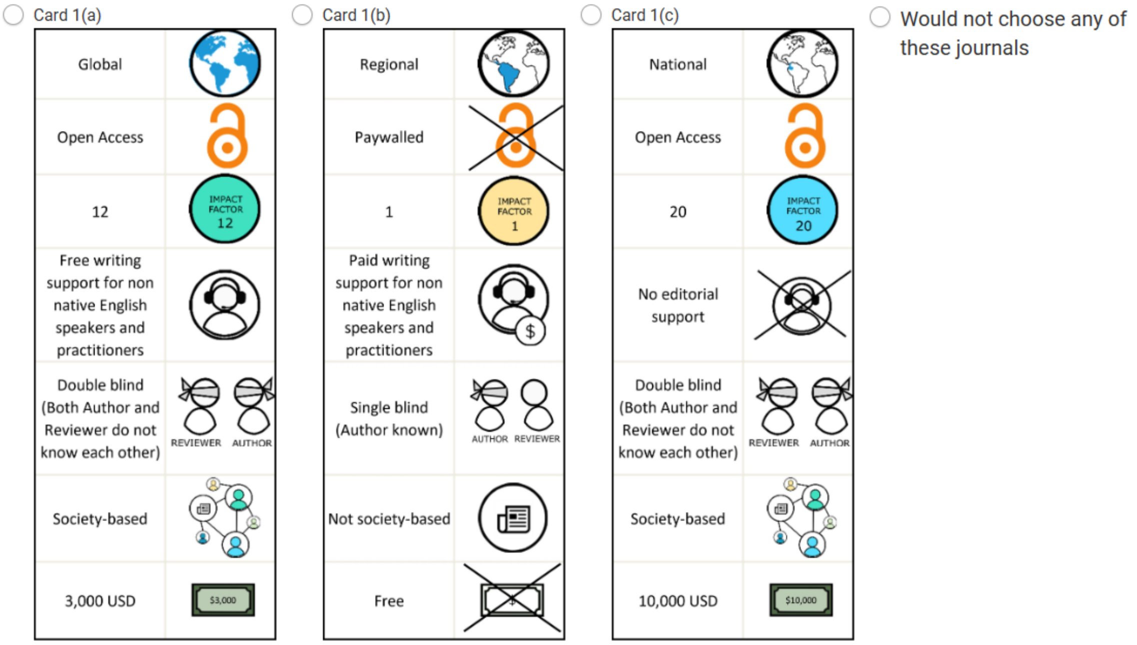


*Example of choice card design*

**Attribute preference**

21. Please rank following attributes from 7 (the most important attribute) to 1 (the least important attribute)

☐ Scope

☐ Access

☐ Impact factor

☐ Editorial support

☐ Review

☐ Society

☐ Cost

22. Have you ignored attributes?

☐ Yes

☐ No

Win a membership to the Society for Conservation Biology!

We are offering all survey respondents the chance to win a membership to the Society for Conservation Biology. We have three 1-year memberships and three 3-year memberships available. If you would like to be considered, please click the link below. If you do not wish to enter the raffle, please select 'Finish survey' to submit your response.

RETRACTED FOR REVIEW

If you have any questions, please speak to RETRACTED FOR REVIEW (RETRACTED FOR REVIEW).

**Appendix 2**. Summary of preliminary surveys at the International Conference of Conservation Biology

To identify important journal attributes to include in the Discrete Choice Experiment, we conducted two surveys during the International Conference of Conservation Biology (2021). Firstly, we hosted a one-hour workshop “Barriers and Opportunities in Academic Publishing” where we divided the 12 attendees (excluding facilitators) into three breakout rooms and each group was asked to discuss two questions, 1) how do you choose where to publish, and 2) what can SCB journals change to better serve the conservation science community? We then compiled the responses from each breakout room. Secondly, we invited conference attendees to complete an online survey where we asked the same two questions along with additional questions, including country of residence and previous experience in publishing conservation research. In total, we received 86 respondents from 33 countries. Seventy-six percent of respondents reported they had previously published in a peer reviewed journal. Topic relevance (30% of respondents), impact factor/reputation (29%), cost (21%), readership (5%), and whether the journal was open access (5%) were the most common factors identified from these surveys. Career type, colleague recommendations, discrimination during the review process, and conservation impact were also identified less frequently.

**Appendix 3**. Information used to invite survey participants. (a) Conservation-related journals used to collect author emails from publications in 2010 and 2020. (b) Conservation-related institutions and organisations whose communication platforms (e.g. newsletters, mailing lists, and social media platforms) were used to advertise the survey more broadly.

**(a) Conservation-related journals**

Aquatic Conservation:
Marine & Freshwater Ecosystems

Biodiversity and Conservation

Biological Conservation

Conservation Biology

Conservation Letters

Conservation Physiology

Conservation Science and Practice

Diversity and Distributions

Ecological Engineering

Environmental Conservation

Global Ecology and Conservation

Human-Wildlife Interactions

Journal for Nature Conservation

Journal of Applied Ecology

Landscape Ecology

Nature Conservation

Oryx

Wildlife Society Bulletin

**(b) Communication channels**

Society for Conservation Biology

Durrell Institute of Conservation and Ecology

Association for Tropical Biology and Conservation

Conservation Leadership Program

Interdisciplinary Center for Conservation Science

The Smith Fellowship

The Rufford Foundation

The Student Conference in Conservation Science (SCCS): Bangalore, Cambridge, Australia

The International Union for Conservation of Nature (IUCN):

- Knowledge Unit
- Species Survival Commission
- Commission on Education & Communication
- Other commissions

National Geographic Explorers

Cambridge Conservation Initiative

**Appendix 4:** Dummy coding used in model estimation to evaluate journal preferences.

| **Variable** | **Type** |
|---|---|
| Journal Scope Global | Dummy |
| Journal Scope Regional | Dummy |
| Journal Scope Local | Dummy* |
| Access Model | Dummy |
| Impact Factor | Score |
| Edit No | Dummy |
| Edit Paid | Dummy |
| Edit Free | Dummy* |
| Peer Review Model | Dummy |
| Society | Dummy |
| Cost | Monetary |

Note: * excluded dummy level

**Appendix 5:** Econometric Model Specifications

Our survey data is used to model a utility function representing respondent preferences over a set of choices. The utility function depends on the attributes of each choice made by respondent *i (i=1,…,N)*. Let $x_{ijt}$ be a $k$×1 vector of attributes presented to respondent $i$ such that they select alternative $j$ ($j$=1,…,$J$) in choice situation $t$ ($t$=1,…,$T$). Assume $U_{ijt}$ is the utility respondent $i$ attains from $x_{ijs}$ such that:

$$U_{ijs} = x'_{ijt}\beta_i + e_{ijt} \quad (1)$$

where $x'_{ijt}\beta_i$ is referred to as the systematic utility component and $e_{ijt}$ is the error term assumed to be extreme value (Gumbel) distributed, independent of $x'_{ijt}$ and uncorrelated across individuals or choices.

LCM Model Specification

LCMs enable estimation of the utility function taking account of respondent heterogeneity. The model specification assumes a fixed (finite) number of classes that respondents are statistically allocated into. The LCM, we employed, modifies equation (1) with the inclusion of an additional subscript, s, that specifies the classes:

$$U_{ijt|s} = x'_{ijt}\beta_i + \varepsilon_{ijt|s} \quad (2)$$

It can then be shown that the probability that respondent $i$, belonging to class $s$, will choose an alternative $j$ from a choice set of $t$ is:

$$Pr_{ijt|s} = \left(\frac{e^{x'_{ijt}\beta_s}}{\sum_{j=1}^{J} e^{x'_{ijt}\beta_s}}\right) \quad (3)$$

As is standard, we employ a MNL to assign respondents $i$ to a specific class $s$:

$$Pr_{is} = \left(\frac{e^{z_i\alpha'_s}}{\sum_{s=1}^{S} e^{z_i\alpha'_s}}\right) \quad (4)$$

where $\alpha_s$ is a vector of parameters to be estimated for each s and $z_i$ is a vector of socio-economic variables explaining class membership. It follows that conditional on a specific class membership that the probability that a $i$ selects $j$ from choice set of $t$ is:

$$Pr_{ij|s} = \prod_{t=1}^{T} Pr_{ijt|s} \quad (5)$$

We estimate the LCM by combining equations (4) and (5) as follows:

$$Pr_{ij} = \sum_{s=1}^{S}\left(\frac{e^{z_i\alpha'_s}}{\sum_{s=1}^{S} e^{z_i\alpha'_s}} \prod_{t=1}^{T}\left(\frac{e^{x'_{ijt}\beta_s}}{\sum_{j=1}^{J} e^{x'_{ijt}\beta_s}}\right)\right) \quad (6)$$

We estimate equation (6) by selecting the number of classes, $s$, and then using maximum likelihood estimation. Model selection in terms of number of classes is conducted using the model log likelihood, the Akaike Information Criterion (AIC), and the Bayesian Information Criterion (BIC).

Mixed Logit (MXL) Model Specification

In the MXL specification (Train, 2009), $\beta_i$ is assumed to be a ($k$×1) vector individual preferences which are independently and identically normal distributed:

$\beta_i = \beta + \Gamma\nu_i$ (7)

where $\beta_i$has a mean $\beta$ and $\Gamma$ is a diagonal matrix including the standard deviation of the random parameter distributions of $\beta_i$ and $\nu_i$ is the unobserved random disturbances.

For our preferred model specification, we have assumed that all parameters follow a normal distribution. To estimate the model in Willingness to Pay space, we follow Balcombe et al. (2010). Our utility function is expressed as follows:

$$U_{ijt} = \beta_1[Cost_{ijt} + \beta_{2,i}Global\,Journal\,Scope_{ij} + \beta_{3,i}Regional\,Journal\,Scope_{ijt} + \beta_{4,i}Access\,Model_{ijt} + \beta_{5,i}Impact\,Factor_{ijt} + \beta_{6,i}No\,Editorial\,Support_{ijt} + \beta_7 Paid\,Editorial\,Support_{ijt} + \beta_8 Peer\,Review_{ijt} + \beta_9 Society_{ijt}] + e_{ijt} \quad (8)$$

where $\beta_{2,i}$ to $\beta_{9,i}$ are WTP parameters for each individual for the set of attributes. This econometric specification does not have a closed form log-likelihood function meaning estimation is implemented using simulated maximum likelihood. Estimation was undertaken using the software NLOGIT Version 6 (Greene, 2016) employing 1000 Halton draws.

To examine the relationship between our WTP estimates and socio-economic data, we follow Xuan et al. (2021) and Blackhall-Miles et al. (2023) and regress individual attribute WTP estimates against the socio-economic data using panel model specification.

**Appendix 6:** Summary of statistical measures of model fit for Multinomial logit (MNL), Latent Class Models (LCM) and Correlated Mixed Logit in Willingness to Pay Space (MXL).

| **Model** | **k[a]** | **LL[b]** | **AIC[c]** | **AIC/N** | **BIC[d]** | **$R^2$** | **S1[e]** | **S2** | **S3** | **S4** |
|---|---|---|---|---|---|---|---|---|---|---|
| MNL | 10 | -14932.3 | 29884.6 | 2.448 | 29958.8 | | | | | |
| LCM2 | 24 | -13229.5 | 26506.9 | 2.171 | 26685.1 | 0.218 | 0.449 | 0.55 | | |
| LCM3 | 38 | -12766.6 | 25609.3 | 2.097 | 25891.3 | 0.245 | 0.234 | 0.309 | 0.457 | |
| LCM4 | 52 | -12471.6 | 25047.2 | 2.051 | 25433.2 | 0.262 | 0.224 | 0.374 | 0.271 | 0.131 |
| MXL | 48 | -11982.7 | 24061.3 | 1.971 | 24417.1 | 0.292 | | | | |

[a]Number of parameters.
[b]Log Likelihood.
[c]Akaike's information criterion.
[d]Bayesian information criterion.
[e]The proportion of respondents by LCM segment (S)

**Appendix 7:** Model Results for the Correlated Mixed Logit in Willingness to Pay Space (MXL). All random parameters Normally Distributed. All coefficients except ASC are Willingness to Pay Estimates. Significance levels: **P* < 0.1, ***P* < 0.05, *** *P*<0.01. ASC – Alternative specific constant.

| **Attributes** | **Coefficients** | | **SE**[a] | **SD**[b] | | **SE** |
|---|---|---|---|---|---|---|
| ASC[c] | -1.266 | *** | 0.077 | | | |
| Global Journal Scope | 75.339 | | 209.321 | 3947.86 | *** | 178.43 |
| Regional Journal Scope | 338.007 | *** | 99.289 | 1536.99 | *** | 113.69 |
| Access Model | 252.100 | ** | 108.186 | 1958.84 | *** | 120.28 |
| Impact Factor | 13.736 | ** | 5.427 | 111.31 | *** | 5.37 |
| No editorial support | -1313.09 | *** | 131.963 | 2293.11 | *** | 185.14 |
| Paid editorial support | -1910.68 | *** | 160.448 | 2555.07 | *** | 212.87 |
| Peer Review Model | 795.326 | *** | 95.985 | 485.82 | *** | 139.45 |
| Society | 130.394 | | 89.459 | 481.12 | *** | 173.97 |

Notes:
a - SE = Standard error
b – SD = Standard deviation
c – ASC is a fixed parameter

**Appendix 8:** Socio-Economic Determinants of Willingness to Pay by attribute using Pooled OLS Fixed and Random Effects Panel Models. Significance levels: *$P$ < 0.1, **$P$ < 0.05, *** $P$<0.01. Excluded level is Regional Journal Scope. GLS Random Effects statistically significant as able to reject null for Bresuch-Pagan and unable to reject null for the Hausmann test.

| | **Pooled OLS** | | **Fixed Effect** | | **GLS Random Effects** | |
|---|---|---|---|---|---|---|
| **Variables** | **Coeff** | **SE[a]** | **Coeff** | **SE[a]** | **Coeff** | **SE[a]** |
| Constant | 500.04*** | 75.73 | 22.54 | 69.49 | 14.59 | 73.89 |
| NoPaper | −3.55 | 2.54 | −5.25* | 2.77 | −4.69* | 2.81 |
| Age | −9.44*** | 1.63 | −9.46** | 1.74 | −9.46** | 1.68 |
| HighInc | 273.16*** | 39.60 | 239.99*** | 42.72 | 252.29*** | 41.57 |
| **Dummies** | | | | | | |
| Global | −379.99*** | 107.27 | | | | |
| Access | −96.33 | 63.43 | | | | |
| IF | −246.83*** | 39.78 | | | | |
| NoEd | −1524.52*** | 71.88 | | | | |
| PaidEd | −2170.39*** | 74.35 | | | | |
| Peer | 535.47*** | 40.70 | | | | |
| Society | −127.32*** | 41.33 | | | | |
| $R^2$ | 0.216 | | 0.32 | | | |
| F Test (Model) | 582.47*** | | 20.07*** | | | |
| F Test (Group Dummies) | | | 8.57*** | | | |
| Regression $\chi^2$ (3) | | | | | 65.34*** | |
| Breusch-Pagan $\chi^2$ (1) | | | | | 1356.25*** | |
| Hausman $\chi^2$ (3) | | | | | 2.64 | |
| N | 8200 | | 8200 | | 8200 | |

Notes: a - SE = Standard error

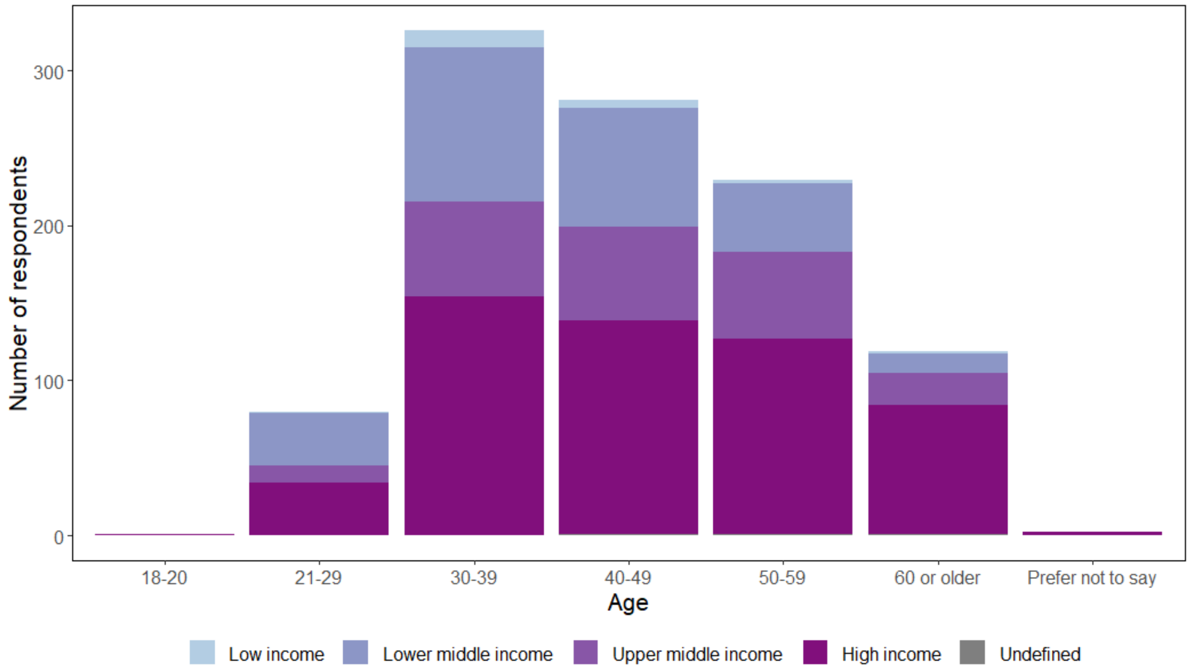


**Appendix 9.** Summary of respondent demographics between age classes and nationality by income group as defined by the World Bank (2022).

**Appendix 10:** Article processing charges (excl. taxes), publishing options, and publisher for each journal included in our study. Article processing charges represent fees for full-length, original research articles in $USD for May 2024.

| Conservation-related journals | Publishing option | Article processing charge ($) | Publisher |
|---|---|---|---|
| Aquatic Conservation: Marine & Freshwater Ecosystems | Hybrid | 4,600 | John Wiley and Sons Ltd |
| Biodiversity and Conservation | Hybrid | 3,790 | Springer Nature |
| Biological Conservation | Hybrid | 3,570 | Elsevier |
| Conservation Biology | Hybrid | Society members 2,592; Non-members 3,240 | John Wiley and Sons Ltd |
| Conservation Letters | Gold open access only | Society members 2,584; Non-members 3,230 | John Wiley and Sons Ltd |
| Conservation Physiology | Gold open access only | Society members 2,195*; Non-members 2,439* | Oxford University Press |
| Conservation Science and Practice | Gold open access only | Society members 1,632; Non-members 2,040 | John Wiley and Sons Ltd |
| Diversity and Distributions | Gold open access only | Society members 3,120; Non-members 2,496 | John Wiley and Sons Ltd |
| Ecological Engineering | Hybrid | 3,650 | Elsevier |
| Environmental Conservation | Hybrid | 3,450 | Cambridge University Press |
| Global Ecology and Conservation | Gold open access only | 2,230 | Elsevier |
| Human-Wildlife Interactions | Gold open access only | 2,125 | Frontiers |
| Journal for Nature Conservation | Hybrid | 2,560 | Elsevier |
| Journal of Applied Ecology | Hybrid | Society members 2,625; Non-members 3,500 | John Wiley and Sons Ltd |
| Landscape Ecology | Gold open access only | 4,390 | Springer Nature |
| Nature Conservation | Hybrid | 2,560 | Elsevier |
| Oryx | Gold open access only | Member rate 2,933; Non-members 3,450 | Cambridge University Press |
| Wildlife Society Bulletin | Gold open access only | Society members 1,980; Non-members 2,200 | John Wiley and Sons Ltd |

*Exchange rate from GBP to USD 1.2710 (May 21st 2024).
